# Denoising High-field Multi-dimensional MRI with Local Complex PCA

**DOI:** 10.1101/606582

**Authors:** Pierre-Louis Bazin, Anneke Alkemade, Wietske van der Zwaag, Matthan Caan, Martijn Mulder, Birte U. Forstmann

## Abstract

Modern high field and ultra high field magnetic resonance imaging (MRI) experiments routinely collect multi-dimensional data with high spatial resolution, whether multi-parametric structural, diffusion or functional MRI. While diffusion and functional imaging have benefited from recent advances in multi-dimensional signal analysis and denoising, structural MRI has remained untouched. In this work, we propose a denoising technique for multi-parametric quantitative MRI, combining a highly popular denoising method from diffusion imaging, over-complete local PCA, with a reconstruction of the complex-valued MR signal in order to define stable estimates of the noise in the decomposition. With this approach, we show signal to noise ratio (SNR) improvements in high resolution MRI without compromising the spatial accuracy or generating spurious perceptual boundaries.

## 1. Introduction

Ultra-high field magnetic resonance (MR) imaging at 7 Tesla and beyond has enabled neuroscientists to probe the human brain *in vivo* beyond the macroscopic scale (Weiskopf et al., 2015). In particular, quantitative MRI techniques (Cercignani et al., 2018) have become more readily available and offer the promise of quantitative information about the underlying microstructure. Unfortunately, as the size of the imaging voxel decreases well below the cubic millimeter, so does the signal to noise ratio (SNR), and the achievable resolution even when imaging a small portion of the brain remains limited, and requires multiple averages at the highest resolutions (Federeau et al., 2018; Fracasso et al., 2016; Stucht et al., 2015).

Multi-parametric quantitative methods acquire multiple images within a single sequence, in order to estimate the underlying quantity (Helms et al. 2008; Metere et al. 2017; Caan et al., 2019). This is comparable to diffusion-weighted MR imaging (DWI) where multiple images with different weighting are acquired. Recent work in diffusion analysis demonstrated that the intrinsic redundancy of the signal across these images can be employed to separate signal from noise with a principal component analysis (PCA) over small patches of the images (Manjon et al., 2013; Veraart et al., 2016). This principle can be transferred to multi-parametric quantitative MRI: in this work we present an extension of the PCA denoising to the recently described MP2RAGEME sequence, which provides estimates of R1 and R2* relaxation rates as well as quantitative susceptibility maps (QSM) in a very compact imaging sequence (Caan et al., 2019).

The MP2RAGEME data comes with additional challenges for the classical PCA approaches. First, the images are complex-valued, comprising of both magnitude and phase needed for the estimation of quantitative MRI parameters. This complex nature of the data needs to be taken into account. Second, they include only a few images compared to the high number of directions acquired in DWI. Interestingly, we can make fairly simple assumptions about the dimensionality of the signal, as it contains primarily R1, R2*, susceptibility and proton density (PD) weighting. Taking these features into account, we were able to effectively denoise high resolution MP2RAGEME data without averaging, and maintain spatial precision. We studied the impact of denoising on the computation of quantitative maps, and its practical impact for reconstructing thin vessels and delineating small, low contrast subcortical nuclei such as the habenula, a notoriously difficult-to-image, small structure in the subcortex.

## 2. Materials and Methods

### 2.1. Data acquisition

Our denoising method focuses on the recently developed MP2RAGEME sequence (Caan et al., 2019) which combines multiple inversions and multiple echoes from a magnetisation-prepared gradient echo sequence (MPRAGE) to simultaneously obtain an estimate of quantitative T1, T2* and susceptibility. The specific protocol of interest is comprised of five different images: a first inversion with T1 weighting, followed by a second inversion with predominantly PD weighting and four echoes with increasing T2* weighting (Fig. 1).

**Figure 1:**
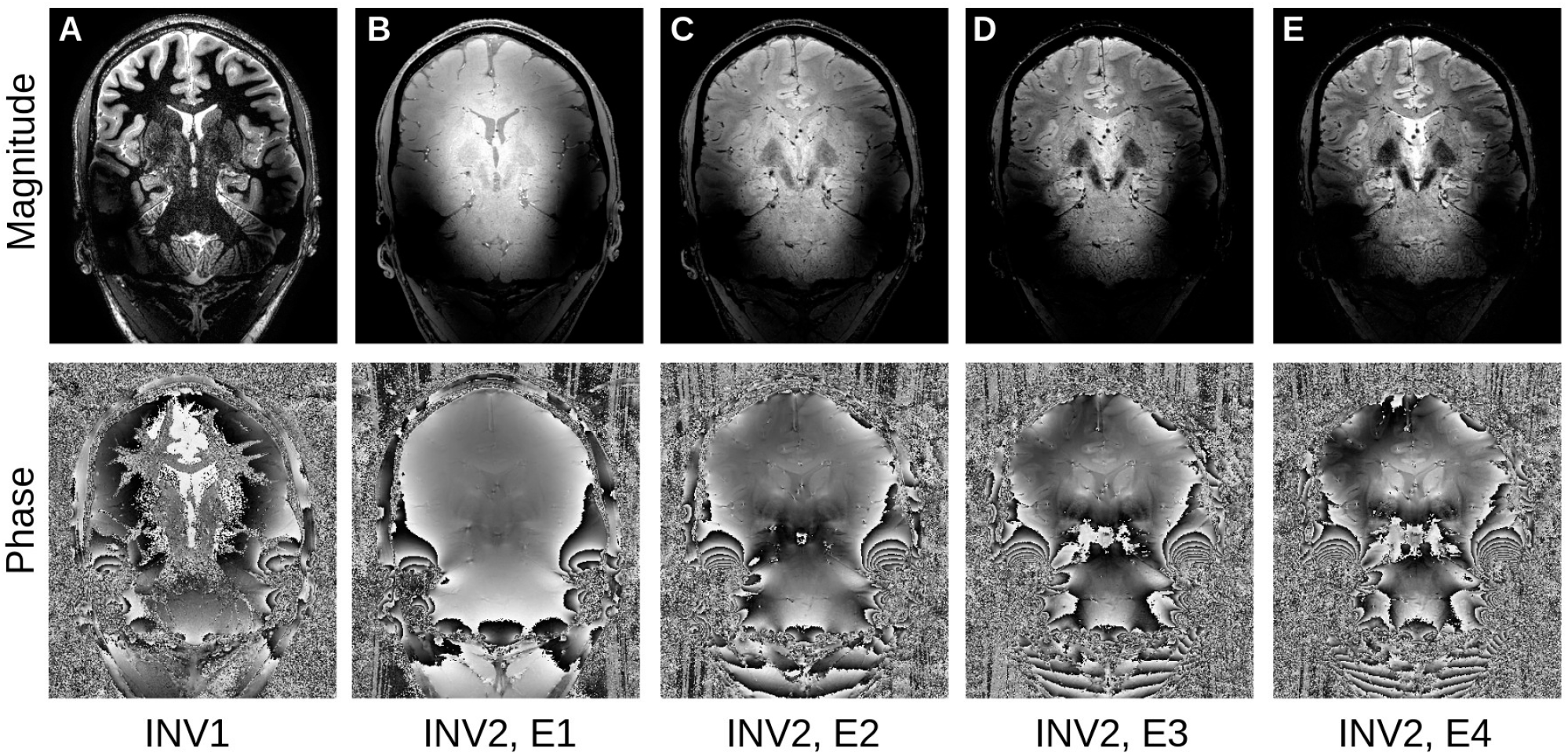
The MP2RAGEME sequence: A: first inversion, B,D,C,E: second inversion, first to fourth echo (top: magnitude, bottom: phase).

For the experiments, we used a high-resolution imaging slab centered on the subcortex with the following parameters: resolution = 0.5mm isotropic, TR = 8.33s, TR_1_ = 8ms, TR_2_ = 32ms, TE_1_ = 4.6ms, TE_2A-D_ = 4.6/12.6/20.6/28.6ms, TI_1_/TI_2_ = 670/3738ms, α_1_/α_2_ = 7/8°. Data was obtained on a 7T system (Philips Achieva, NL) with a 32 channel rf-coil (Nova Medical Inc, USA) for five healthy human subjects as part of an ongoing atlasing study approved by the Ethics Review Board of the Faculty of Social and Behavioral Sciences, University of Amsterdam, The Netherlands (approval number: 2016-DP-6897). All subjects provided written informed consent for the study.

### 2.2. Complex signal reconstruction

One of the main drawbacks of existing local PCA methods for denoising is that they work on magnitude images, which follow Rician distributions. A simple correction proposed in (Baselice et al., 2009; Eichner et al., 2016) for removing the Rician bias in DWI averaging is to revert to the complex signal, taking into account the phase information. Phase contains additional variations due to non-local effects of air cavities around the brain, which bring severe ringing artifacts in the reconstructed data (Fig.2A). In (Eichner et al., 2016), the global phase information is removed with a total variation method, which generally respects the location of phase wraps. However, we found that this approach is not effective in regions where the phase wraps have high frequency, and there are residual phase artifacts in the local phase. While these have little consequences for their averaging application, they provide systematic variations for the PCA decomposition, which is undesirable here. We therefore start with a full phase unwrapping (Abdul-Rahman et al., 2005) followed by a total variation smoothing (Chambolle, 2004) of the unwrapped phase (Fig.2B). The residual phase variations are used as local phase, and combined with the magnitude to reconstruct real and imaginary parts of the complex signal (Fig.2C,D). Note that here, unlike (Eichner et al., 2016), we do not discard the imaginary part of the signal as it contains valuable information about the noise and residual anatomical information.

**Figure 2:**
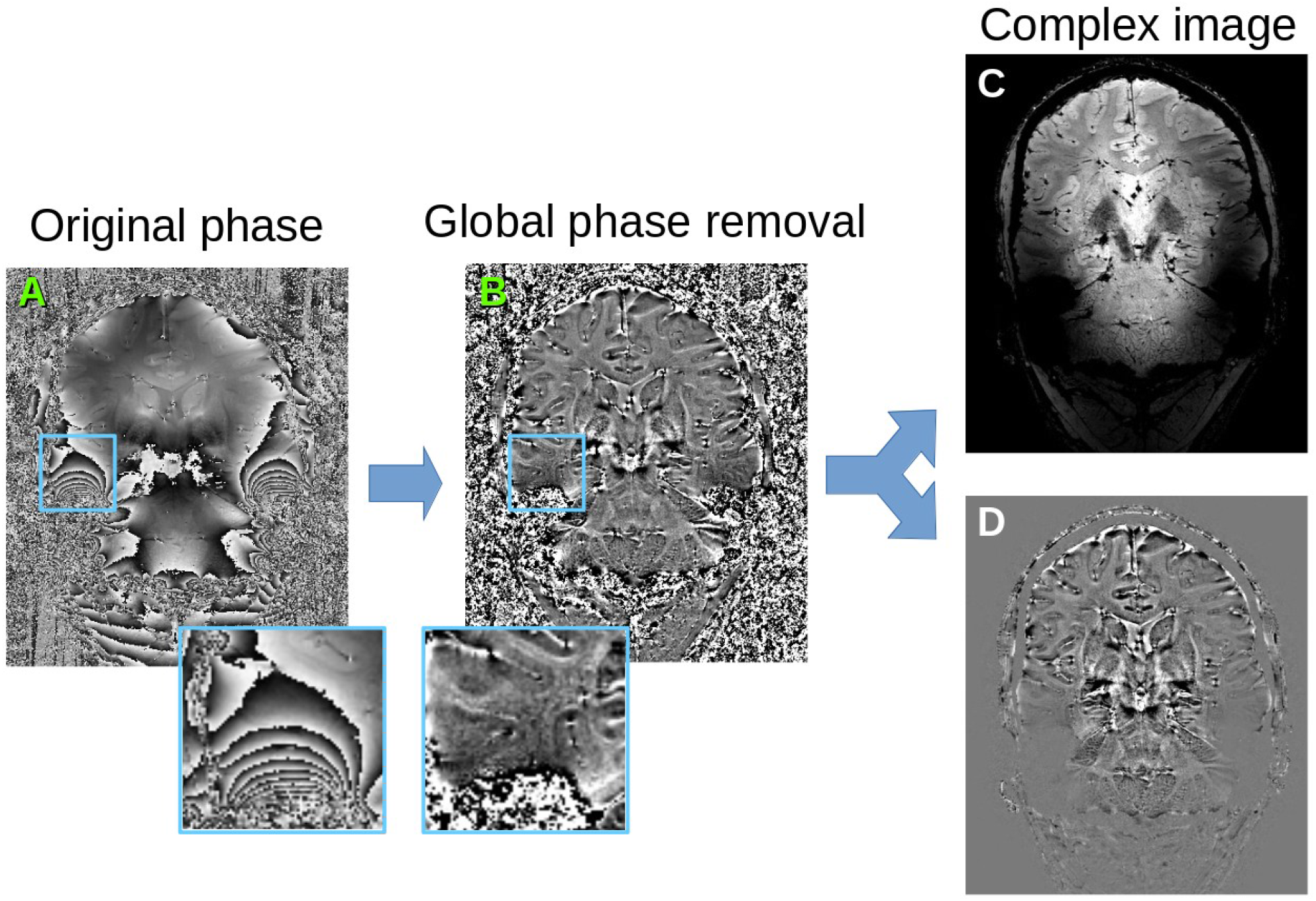
Phase preprocessing. A: original phase map, B: after unwrapping and global phase removal (estimating global phase variations with TV-smoothing and keeping only the residual phase), C: reconstructed real signal, D: reconstructed imaginary signal.

### 2.3. Local PCA of complex signal

The complex signal, now comprising ten image dimensions all similar in nature, is then processed following the local overcomplete PCA approach of (Manjón et al., 2013). In short, the images are cut into small, overlapping patches of *NxNxN* voxels, and the *M* contrasts combined into a *N^3^×M* matrix. The average patch value per contrast is subtracted, and the matrix decomposed via singular value decomposition (SVD) to yield the eigenvectors and associated singular values (square roots of the eigenvalues) of the covariance matrix across the patch. In this work, we used patches of size *N* = 4, and *M* = 10.

The overlapping patches are combined following the technique of (Manjón et al., 2013), weighting each patch by the number of kept eigenvectors 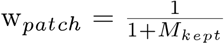. Once recombined, the eigenvectors rapidly change from a rich information content, encoding boundaries, to pure noise (Fig. 3). However, the decision boundary between actual signal and noise is variable across the brain, due to the presence of different tissue types, multiple tissue boundaries, etc. In order to infer for each patch the number of eigenvectors to keep, we thus need to first quantify the expected noise distribution over the SVD.

**Figure 3:**
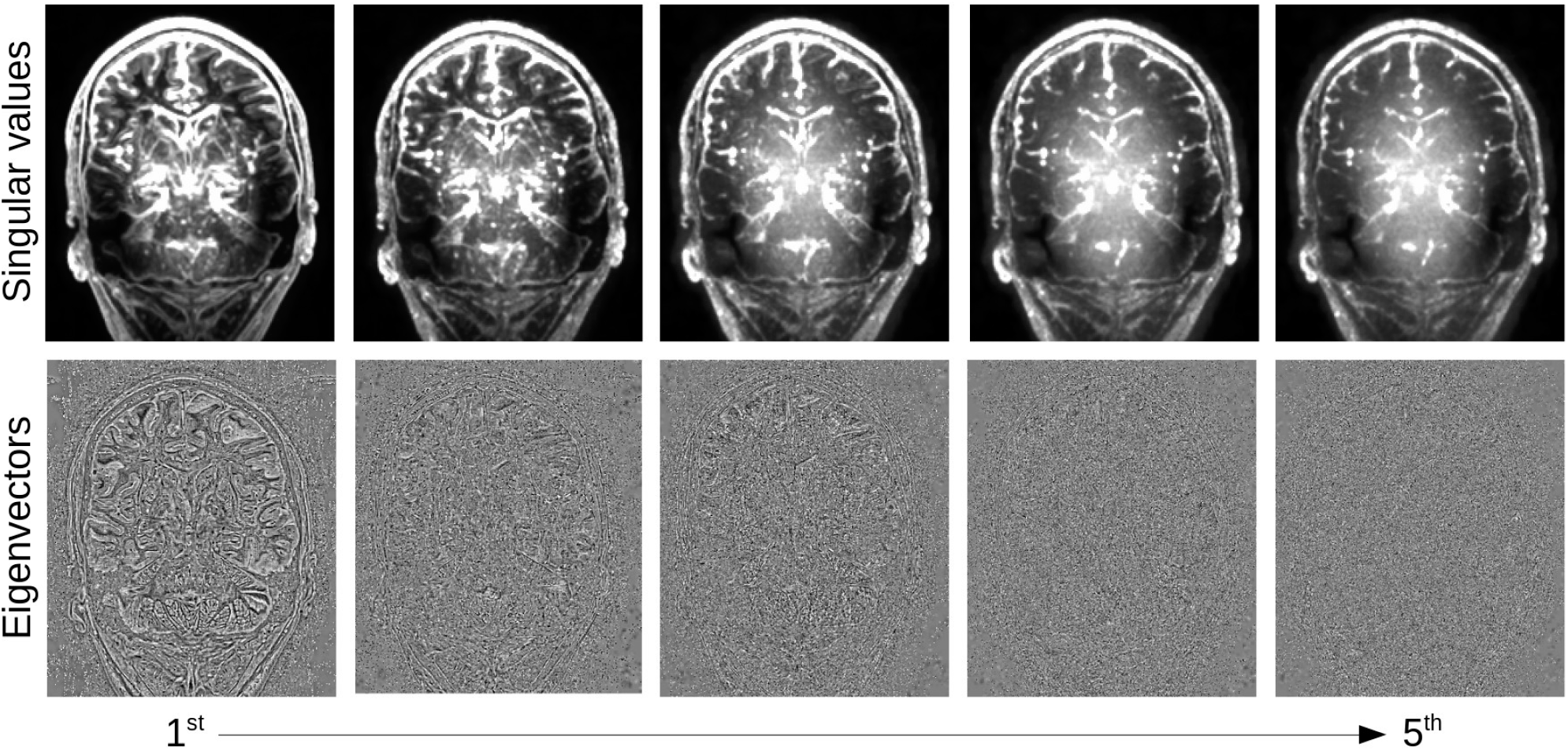
Local PCA decomposition: first five eigenvectors (top) and singular values (bottom) from highest to lowest.

### 2.4. Estimating the noise

Noise estimation in advanced MRI is challenging: variations in coil sensitivity, non-local susceptibility effects and dependencies to the B0 field as well as various acquisition techniques will affect the signal and the noise differently in different regions. The original local PCA denoising methods of (Manjón et al., 2013) used elaborate estimates of the magnitude image noise, taking into account its Rician properties. A recent extension of the work uses random matrix theory to model the expected distribution of random noise eigenvalues and determine its threshold (Veraart et al., 2016). While this approach is theoretically grounded, it may be sensitive to disturbances when applied to data with a limited number of eigenvalues. In Fig. 4, we simulated small 4×4×4 voxel patches with 10 dimensions (as in the MP2RAGEME sequence) and 3 non-zero eigenvalues. While the random matrix theory method performs well in non-interpolated data, its performance degrades strongly when the noise becomes correlated across voxels from image interpolation.

**Figure 4:**
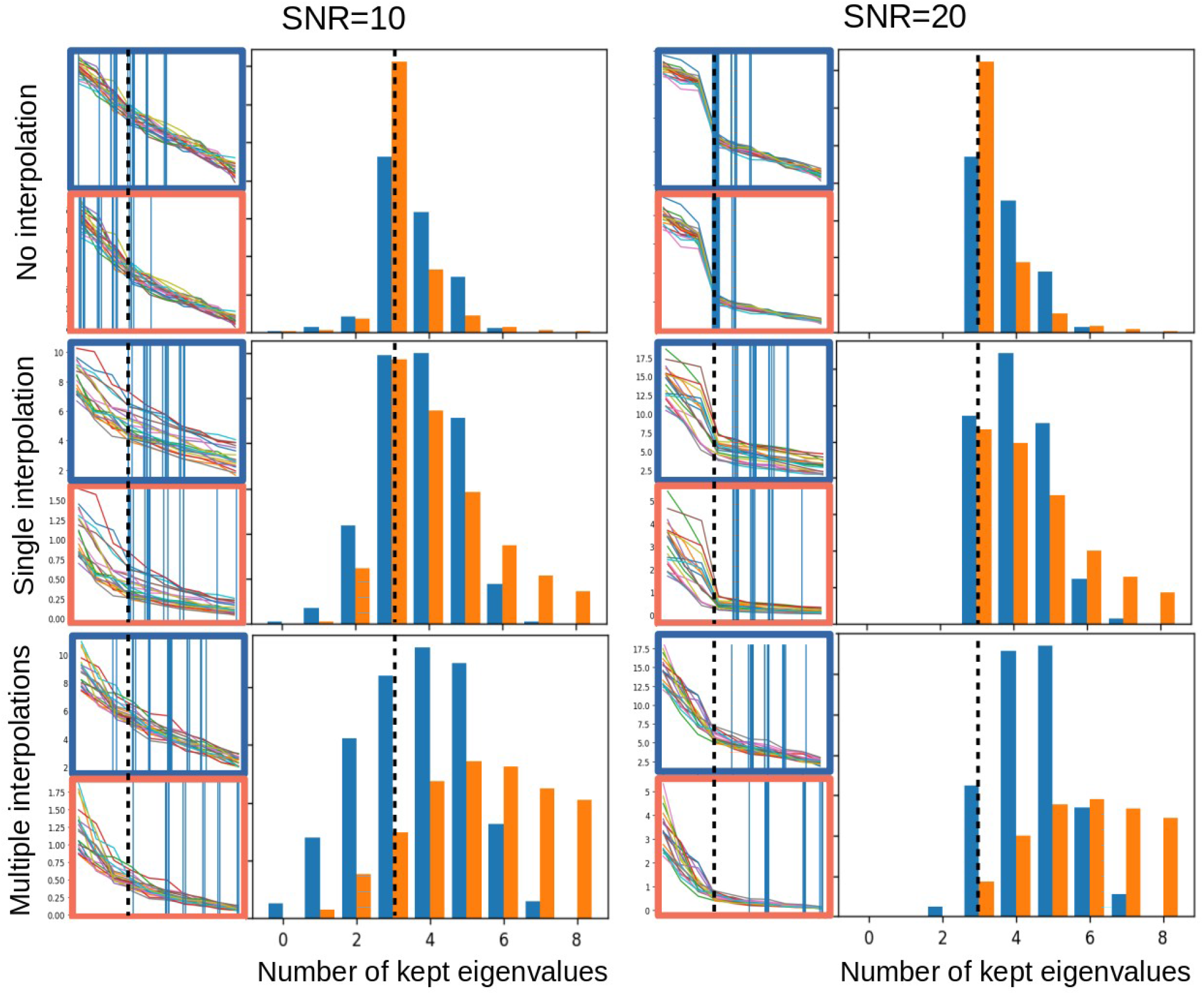
Noise properties of the PCA thresholding: estimation of the threshold to denoise a 4×4×4 voxel patch with 10 dimensions and 3 non-zero eigenvalues, with added Gaussian noise and a SNR of 10 (left) or 20 (right), with no interpolation (top), a single interpolation of all 10 dimensions (middle), or multiple linear interpolations for each of the 10 dimensions (bottom). All interpolations are linear interpolations by a uniformly distributed offset of up to a half voxel. In each subfigure, the left side shows 20 eigen- and singular value examples with the corresponding threshold as a vertical bar, for the random matrix approach (in red) and the linear fitting approach (in blue). The right side shows the histogram of thresholds estimated over 10,000 simulated noise patterns. Dotted lines indicate the ideal number of eigenvalues to keep.

In this work, we propose a more robust approach based on two properties of the complex signal: first, that the noise is locally Gaussian, and second that the spread of singular values for Gaussian noise can be reasonably well approximated by a straight line, at least when the number of dimensions in the decomposition is small. This property is retained by interpolation, which makes it possible to perform denoising after image registration, for instance when motion correction is needed as in DWI or fMRI, or after fat navigator-based motion correction in structural MRI (Gallichan and Marques, 2017).

Our algorithm to estimate the noise level proceeds as follows: first, a line is fitted to the *M*/2 lowest singular values of the patch decomposition using linear least squares (here, *M*/2=5). Then, every singular value above a factor of *α* above the expected noise level given by the fitted line is kept, while the others are removed (in this work, we used *α* = 5%). Thus, the main requirements of our method are that: 1) local signal variations across contrasts in each individual patch are Gaussian-distributed, and 2) the intrinsic dimension of the data is lower than half of the number of acquired images. The patches are then reconstructed and averaged across the image, the complex images separated into magnitude and phase, and finally the discarded global phase variations are reintroduced and wrapped, to obtain data as similar as possible to the original input. In addition, a map of the number of kept eigenvectors and of the noise fitting residuals weighted by patch are computed for quality control (see Fig. 5). The complete denoising algorithm is available in Open Source as part of the IMCN Toolkit (https://www.github.com/imcn-uva/imcn-imaging/) and the Nighres library (Huntenburg et al., 2018; https://www.github.com/nighres/nighres/).

**Figure 5:**
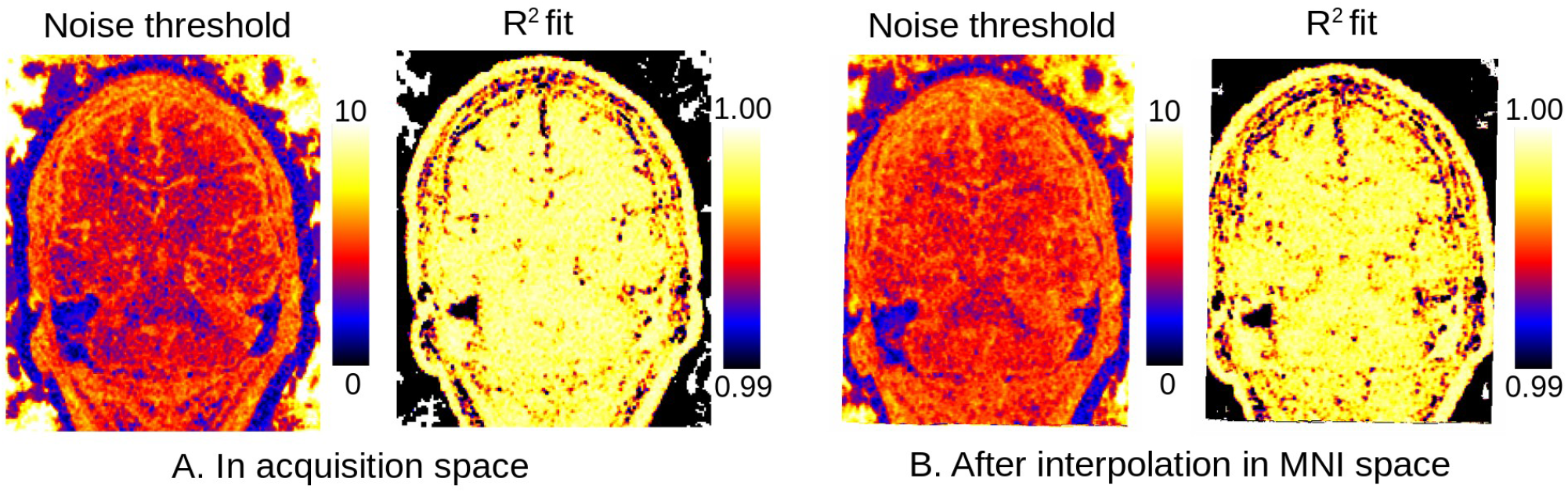
Maps of estimated noise threshold from actual data: number of kept eigenvectors (left) and R^2^ goodness of fit of the linear fit of the lowest singular values of the PCA (right) for data in original acquisition space (A) and linearly co-registered into MNI space (B).

### 2.5. Quantitative mapping

The main interest of quantitative MR mapping techniques such as the MP2RAGEME sequence is to obtain estimates of the MR parameters of T1, T2* relaxation times and susceptibility χ from the measured images. In order to do so, we used the look-up table method of (Marques et al., 2010) to generate T1 estimates, a simple regression in log domain to obtain T2* (Miller and Joseph, 1993) and the TGV-QSM reconstruction algorithm of (Langkammer et al., 2015) to create quantitative susceptibility maps (QSM), following the approach of (Caan et al., 2019) to estimate QSM for each of the three last echoes of the second inversion and take the median. As a byproduct, the T1 estimation also generates bias-corrected T1-weighted images, and the T2* fitting produces an S0 baseline image devoid of T2* effects, which will be used in some of the processing. T1 and T2* mapping methods are included in the Nighres library, and the TGV-QSM algorithm is also freely available (http://www.neuroimaging.at/pages/qsm.php).

### 2.6. Manual delineations

In order to obtain structure-specific measures of quality and to evaluate the practical benefits of the method for segmentation purposes, we performed a manual segmentation study. To evaluate the benefits of the denoising, we selected the habenula, a small structure ventral to the thalamus, which together with the pineal gland and stria medullaris forms the epithalamus (Mai et al, 2016; Hikosaka, 2010; Strotmann et al., 2013). Since we cannot distinguish between the medial and lateral habenula on MRI, both were included in our delineations. Given its size, complex shape, and heterogeneous composition, the habenula requires high resolution images and detailed anatomical knowledge to be reliably delineated on MRI.

In this study, the habenula was delineated by two raters on the reconstructed quantitative T1 maps obtained with or without denoising. The consensus mask (regions labeled by all raters on all images) were used to measure noise properties, while the overlap of the masks was used to measure inter-rater reliability.

### 2.7. Co-registration to standard space

Previously proposed methods for local PCA denoising have shown decreased performance when handling interpolated data due to the changes in the noise distribution (Manjón et al., 2013; Veraart et al., 2016). To test the performance of our noise estimation approach, we co-registered our data to standard space by aligning it first with a whole brain image of the same subject and then to the MNI152 template at 0.5 mm resolution. Both registrations used the first inversion of the MP2RAGEME sequence, and were performed with linear registration in ANTS (Avants et al., 2008). Phase images were unwrapped before transformation to avoid interpolating across phase wraps, and quantitative maps were computed before and after denoising in standard space.

### 2.8. Region labeling

To evaluate the signal improvement across a variety of structures, we used manual delineations of the striatum, globus pallidus internal and external segment, subthalamic nucleus, substantia nigra and red nucleus obtained from lower resolution MP2RAGEME images (0.64×0.64×0.7 mm resolution) with higher SNR. The delineations were performed independently by two raters and the conjunction between them was used as the structure mask. The masks were co-registered linearly to the high resolution slab based on their scanner coordinates followed by a rigid registration in ANTS. The intensities of the quantitative maps inside each region were averaged and used to compute signal-to-noise ratio statistics of the original images and reconstructed quantitative maps before and after denoising in original space.

### 2.9. Vascular reconstructions

Finally, we tested the capabilities of the denoising to maintain fine details by segmenting the vasculature with the method of (Huck et al., 2019) on R2* quantitative maps. The algorithm uses a spatial vessel filter followed by global diffusion, and is particularly sensitive to small vessels (Bazin et al., 2016). The filter was run with standard parameters on both the original and denoised R2* maps after skull stripping using the S0 image and T1 map obtained during quantitative mapping (Bazin et al., 2014), and the result was overlaid on the bias-corrected T1-weighted image for orientation.

## 3. Results

### 3.1. Impact on quantitative maps

As shown in Fig. 6, the LCPCA denoising strongly impacts the estimation of quantitative signals, as the SNR gains of multiple images are combined. Visually, we found a strong gain in particular for R2* mapping, which is quite noise-sensitive with the low number of echo times of the MP2RAGEME sequence. While the improvements are subtle at the whole brain scale, they are very clear when focusing on small regions, particularly in the subcortex where the original SNR is low.

**Figure 6:**
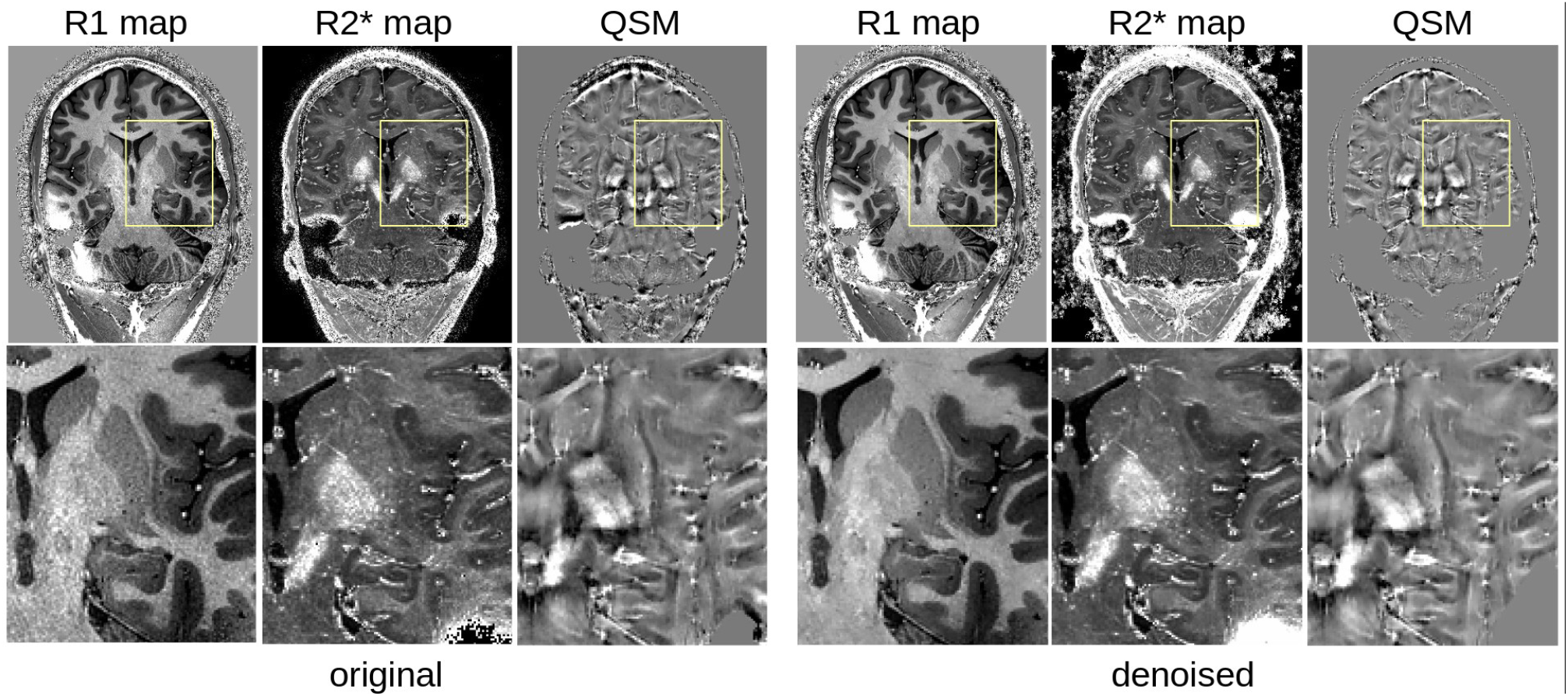
Comparison of quantitative MRI reconstruction: left, from the original MP2RAGEME images; right from the denoised result.

### 3.2. SNR measures

The denoising systematically improved the SNR in all structures, although not identically across regions and contrasts (Fig. 7). On individual MP2RAGEME images, the improvement was modest, while denoising had a stronger cumulative impact on quantitative map estimation, especially for R1. While R2* appears visually improved, the SNR measures were more similar partly due to the heterogeneity of R2* values in many anatomical structures. QSM is the least affected, both visually and quantitatively, which is expected as the quantitative susceptibility reconstruction algorithm includes its own spatial regularization method (in this case, total generalized variation (TGV)). Still, some more heterogeneous regions of the subcortex appear smoother after denoising. Interestingly, the highest gains were obtained for the smaller structures such as substantia nigra, subthalamic nucleus and red nucleus.

**Figure 7:**
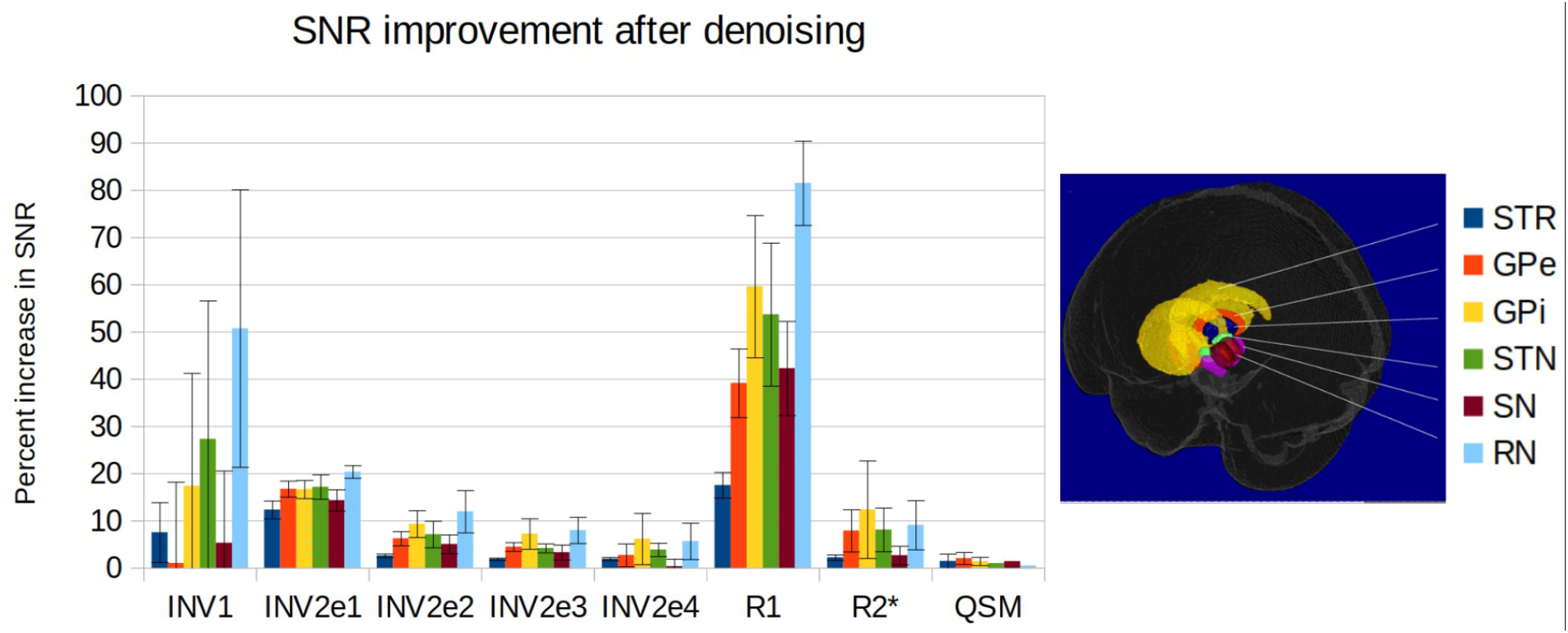
SNR improvement for the delineated structures and the different contrasts (from left to right: first inversion, second inversion echo 1 to 4, estimated R1 map, estimated R2* map, estimated QSM; mean and standard deviation across subjects).

### 3.3. Impact on manual delineations

Manual delineations of the habenula are challenging, and inter-rater agreement is low. Our denoising improved the consensus (Fig. 8), but not to the point of an acceptable level of reproducibility for measuring anatomical quantities such as volume or shape.

**Figure 8:**
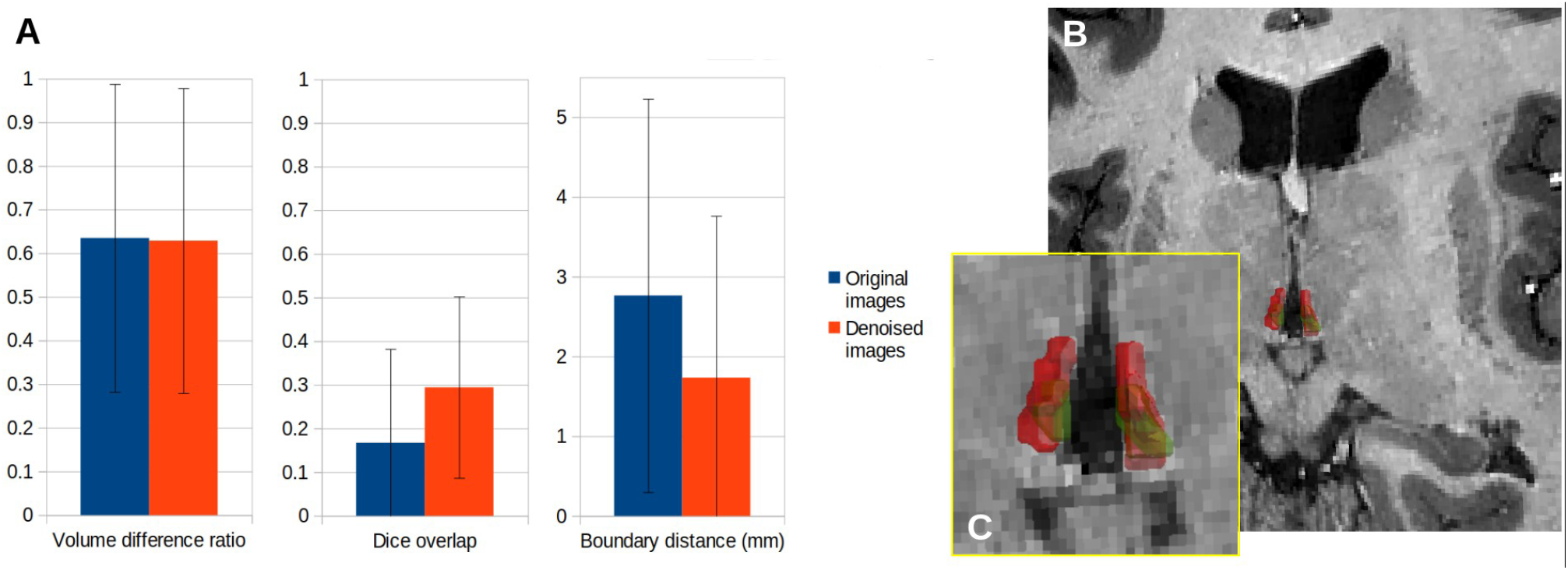
Comparison of manual delineations of the lateral habenula with or without denoising. A: Inter-rater agreement between delineations, B: 3D rendering of the lateral habenula in one subject, as delineated by each rater (red and green outline, respectively), C: Zoomed-in rendering.

### 3.4. Effects of interpolation

When performing the denoising step after image co-registration to a standard space and interpolation, we observe only small differences in the LCPCA algorithm behavior (Fig. 5). The interpolation procedure tends to slightly increase the dimension of the kept signal, probably due to the mixing of signals across voxels. However, the proposed dimensionality estimation procedure appears largely robust to interpolation.

### 3.5. Impact on vascular reconstruction

Brain vasculature is very difficult to image, as most of the vessels have sub-millimeter resolution even at the surface of the brain. Because most methods use local information to identify vessels, they are very sensitive to noise, and prone to large amounts of false detections. Fig. 9 shows the benefits of the proposed denoising in separating vessels from noise, while preserving the fine details of the vasculature.

**Figure 9:**
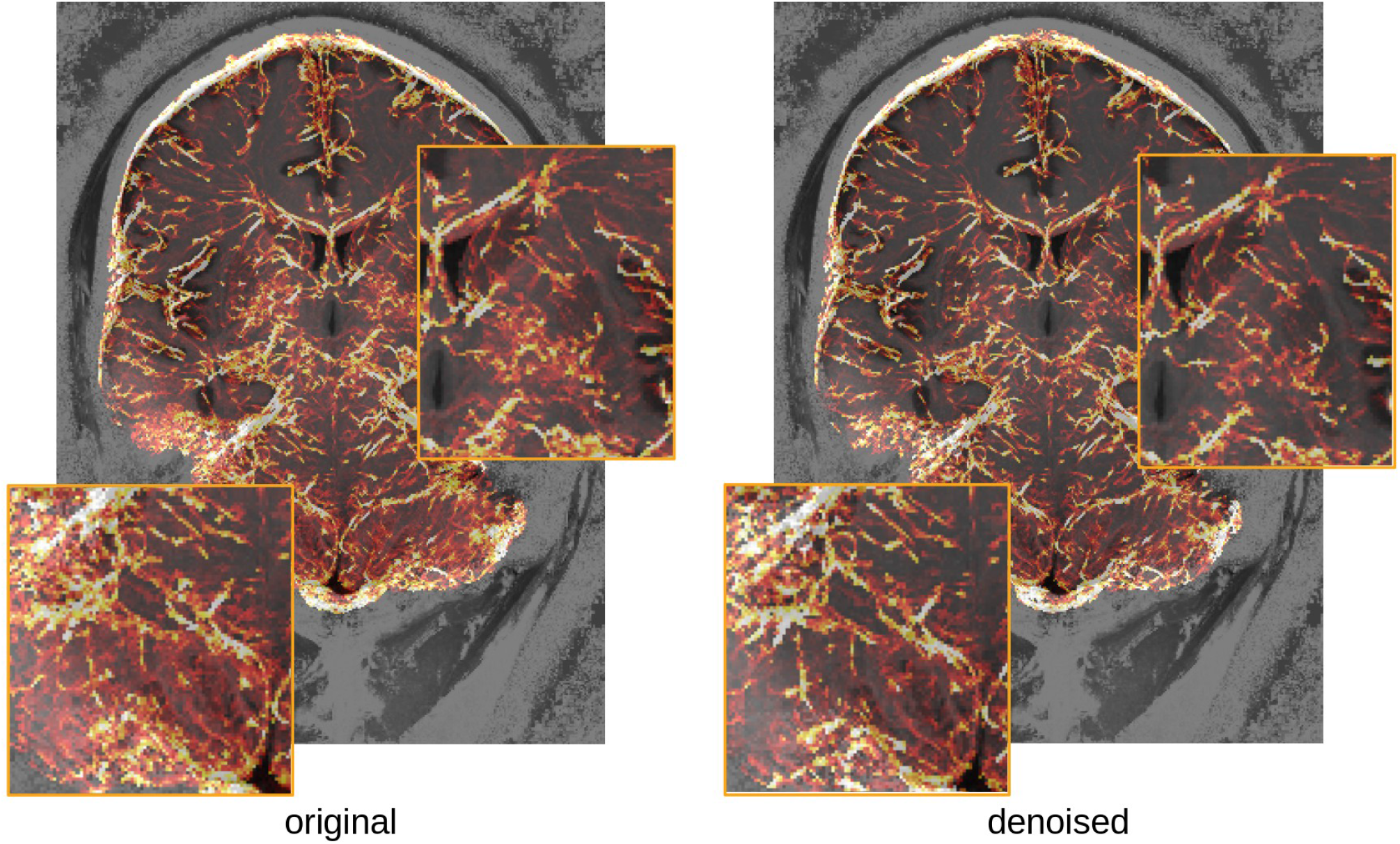
Comparison of vasculature reconstruction with or without denoising (maximum intensity projection over 15mm of the estimated vessel probability map (overlaid on the T1-weighted image for the subject, for orientation). Insets show regions of the cerebellum (left) and subcortex (right) where lower SNR artificially increases the detected vascular density in the original image.

## 4. Discussion

In this work we presented a new local PCA denoising technique, using the full complex signal acquired to correct acquisitions with multiple image contrasts such as DWI or quantitative MRI sequences. Even with a sequence like MP2RAGEME, where only five different images are used in order to recover T1, T2* and QSM, the proposed approach was able to effectively reduce the noise without introducing blurring artifacts. The homogeneity of structures compared to their boundary was improved over the basal ganglia structures, and delineation of small brain structures such as the habenula was shown to be more reproducible in high resolution images with high levels of imaging noise.

By using both magnitude and phase information, the proposed method captures the entire noise distribution, and stays in the statistically simpler domain of complex Gaussian perturbations. However, obtaining high quality phase images is not trivial for all scanners and imaging sequences and can be a limitation also in retrospective processing, as the phase is commonly discarded. Note also that certain acceleration techniques such as partial Fourier encoding may sacrifice the phase signal estimation. While there is no conceptual requirement to use both phase and magnitude in the denoising, lowering the number of dimensions reduces the applicability of the method: in the case of the MP2RAGEME using only five dimensions in order to separate four-dimensional signals from noise is challenging, and the linear fit of the noise to the last half of the local PCA eigenvalues becomes unreliable. Yet, preliminary experiments with other relaxometry methods such as multiparameter mapping (Weiskopf et al., 2013), which acquire between 14 and 20 images, indicated that the separation of signal and noise based on magnitude alone was possible (unpublished data).

The main requirements of the noise estimation method are that: 1) local signal variations across contrasts in each individual patch are Gaussian-distributed, and 2) the intrinsic dimension of the data is generally lower than half of the number of acquired images. The first requirement is easily met in small patches, regardless of the type of data acquired. The second requirement depends on the type of MR sequence and tissue properties under consideration, but can be met simply by running twice the same sequence, as is commonly done for increasing SNR by classical averaging. Note also that in the case of a single contrast PCA approach reduces to simple averaging with no added value.

The main interest of the PCA-based approaches for denoising high resolution images is that unlike most other methods, they do not impose spatial regularity but rather enforce regularity across contrasts, preserving small anatomical details without blurring or inducing artificial boundaries. In addition, while the impact of noise removal on conventional MR images remains subtle, it offers the option to push MR sequences beyond the usually accepted limits of thermal noise as long as enough signal remains to be reliably detected. As ultra-high field advances toward higher and higher resolutions, such denoising methods may become essential part of the imaging protocol for multicontrast anatomical imaging in the same way they have become a standard tool for advanced DWI pre-processing (Veraart et al., 2016).

Finally, the proposed method is generally applicable to other relaxometry sequences such as multi-echo GRE, multi-echo MPRAGE or multi-parametric maps (MPM), as well as DWI or (multi-echo) fMRI. In some cases, e.g. T2* relaxometry or multi-echo fMRI, it is important to note that the known relationship between echoes could also be used to further discriminate signal from noise (Kundu et al., 2017). The robustness of the proposed noise threshold estimation technique to interpolation makes it also interesting for other MR imaging protocols that include multiple images with different contrasts, even if acquired sequentially or even at different resolutions, as long as the ratio of total number of images to actual number of measured contrast mechanisms is sufficient to estimate noise properties.

## 5. Conclusions

Here we presented a new local PCA method to denoise high resolution multi-parametric quantitative MRI data. Combining magnitude and phase data, we could differentiate between signal- and noise-induced variations with a simple model that is robust to interpolation. The resulting quantitative images are automatically regularized and additional anatomical detail is visible in low contrast regions. The denoising software is openly available as part of the IMCN Toolkit (https://www.github.com/imcn-uva/imcn-imaging/) and the Nighres library (https://www.github.com/nighres/nighres/). The proposed method can be extended to denoise other MR imaging sequences with similar properties, namely partially redundant contrasts and low intrinsic dimensionality. While the method is already efficient to denoise MR images acquired with cutting edge methods at the lower limits of SNR, we hope they may help further to push MR imaging toward acquiring even more challenging data where the noise may visually dominate but significant amounts of signal are still available.

## 6. Author Contributions

The method was developed by PB; data was acquired and pre-processed by MM, WZ, and MC; validation experiments were performed by PB and AA; manuscript preparation and editing was done by PB, AA, WZ, MC, MM, BF.

## 8. Acknowledgments

We would like to thank Josephine Groot for her help in recruiting and scanning subjects and Jill van Dorp and Judith Blankenagel for their help in anatomical delineations.

## 10. Data Availability Statement

All of the data from this study cannot be shared publicly because of current GDPR privacy regulations. However, statistical tables and a representative data set for a single subject can be accessed openly here: https://uvaauas.figshare.com/projects/Denoising_High-field_Multi-dimensional_MRI_with_Local_Complex_PCA/61832.

